# RNA compensation: A positive feedback insulation strategy for RNA-based networks

**DOI:** 10.1101/2021.10.26.465912

**Authors:** Baiyang Liu, Christian Cuba Samaniego, Matthew Bennett, James Chappell, Elisa Franco

## Abstract

The lack of signalling modularity of biomolecular systems poses major challenges toward engineering complex networks. An important problem is posed by the consumption of signaling molecules upon circuit interconnection, which makes it possible to control a downstream circuit but compromises the performance of the upstream circuit. This issue has been previously addressed with insulation strategies including high-gain negative feedback and phosphorylation-dephosphorylation reaction cycle. In this paper, we focus on RNA-based circuits and propose a new positive-feedback insulation strategy to mitigate signal consumption. An RNA input is added in tandem with transcription output to compensate the RNA consumption, leading to concentration robustness of the input RNA molecule regardless of the amount of downstream modules. We term this strategy RNA compensation, and it can be applied to systems that have a stringent input-output gain, such as Small Transcription Activating RNAs (STARs). Our analysis shows that RNA compensation not only eliminates the signaling consumption in individual STAR-based regulators, but also improves the composability of STAR cascades and the modularity of RNA bistable systems.

## 1 Introduction

A long-standing goal in engineering biology is the synthesis of artificial molecular networks with complex dynamics [1, 2]. An important challenge toward this goal is posed by the poor modularity of molecular signalling, that makes it difficult to compose molecular devices [3, 4, 5, 6, 7]. In engineering, modularity means that the inputs and outputs of a component can be coupled to other components without causing a departure from the behavior of each part in isolation. In biology, molecular inputs and outputs interact via binding events that always affect the level of free inputs, outputs, and of energy-storing molecules, introducing context-dependent alterations of the components involved. These effects have been described by concepts such as retroactivity and resource limitation, which have spurred the development of methods to improve modularity by introducing negative feedback and high-gain mechanisms [5, 8, 9, 10, 11, 12]. As the toolkit of synthetic biology is rapidly expanding, new design methods making it possible to build modular molecular systems are highly sought-after [13].

Modularity is becoming critical in the emerging class of RNA-based genetic circuits, which offer many potential advantages like rapid programmability, portability across hosts, and low metabolic burden[14]. While recent work has demonstrated dynamic systems that use exclusively RNA molecules as regulators propagating information[2, 15, 16, 17, 18, 19, 20, 21, 22], the composition of RNA circuits poses unique challenges when compared to more established protein-based circuits. For example, Small Transcription Activating RNAs (STARs) are a synthetic class of small RNA regulator able to activate transcription. This regulator is composed of a target RNA placed upstream of a gene to be regulated that once transcribed, folds into an intrinsic terminator hairpin that prevents transcription elongation of the downstream gene. Activation is achieved through transcription of a STAR that is designed to interact with the target RNA, preventing hairpin formation, and thus allowing paused elongation complexes to recover and to proceed into the downstream gene (Fig 1A). Once a STAR regulator is fully bound to its target RNA, this interaction can be considered irreversible at 37°C because of extensive RNA:RNA interactions (>60 bp) [15] with high predicted free energy of binding (Δ*G* ≈ −70*kcal/mol*) and small dissociation constants (*K_D_* ≈ 6 × 10^−53^*mol/L*). Thus, each successful STAR-mediated activation event results in consumption of a STAR molecule; similarly, RNA-based repressors operate by depleting RNA regulators. This is in contrast with the behavior of transcription factors in protein-based circuits, in which proteins can bind and unbind dynamically from their targets and are not permanently depleted. The consumption of RNA regulators leads to a substantial perturbation of the desired levels of regulators, and especially limits the scalability of STAR circuits.

**Figure 1:**
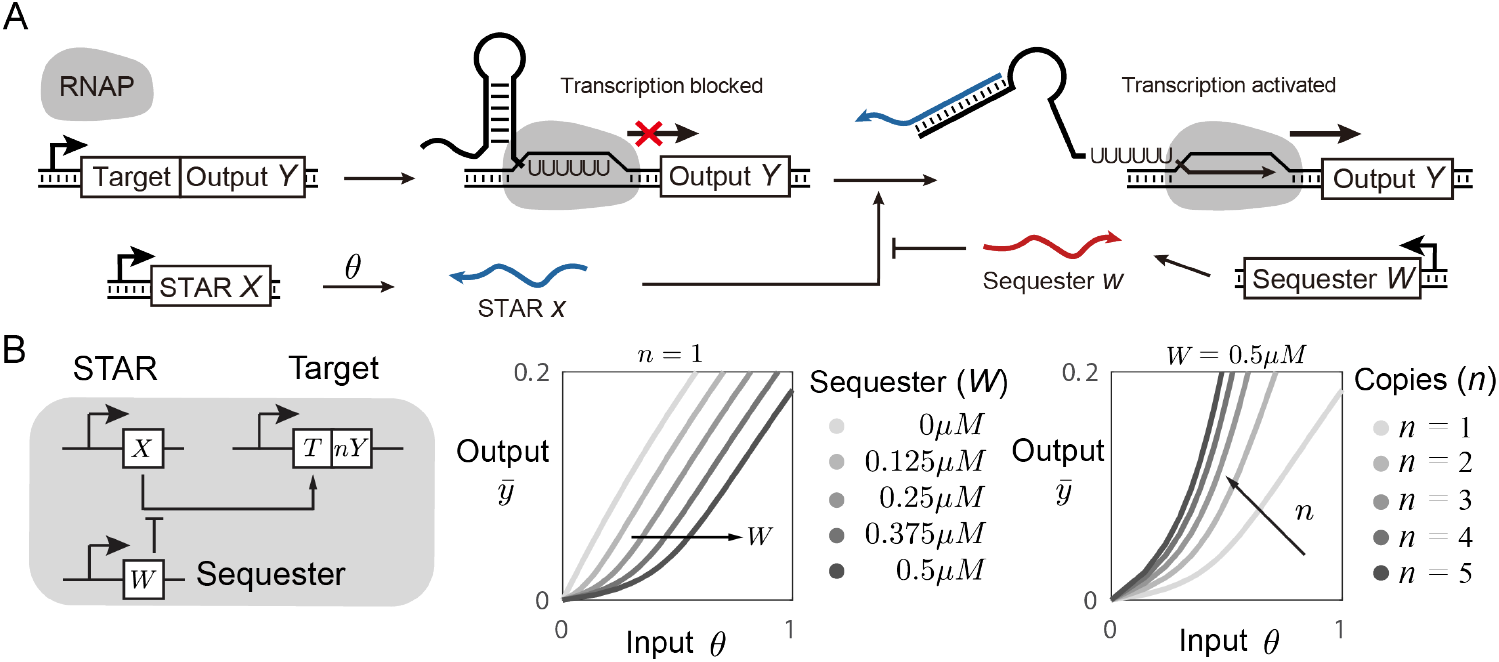
Tunable STAR regulators. (A) A schematic of the STAR-Target RNA regulatory system. A target RNA (T) that encodes a terminator is placed upstream of an output gene (Y). A constitutive promoter initiates transcription of the target RNA, which by default, folds into a terminator to pause and then terminate transcription elongation by RNA polymerase (RNAP). In the presence of the STAR (X) the target RNA folds into an alternative antiterminated structure, which allows transcription of the output RNA. Activation by STAR can be inhibited by expression of an RNA sequester (W) that forms duplex interactions with the STAR. (B) Left panel - the schematic of STAR with sequestration and output amplification. Middle and right panel - simulation results showing that the presence of RNA sequester and tandem outputs allow non-linear and tunable response in the STAR system. The input θ annotates the transcription rate of STAR X. n annotates the repeat number of output Y.

The fundamental mechanistic differences in how regulation is achieved by RNA and protein-based regulators pose unique challenges and opportunities to the design of composable circuits. For example, while potentially problematic, the precise nature of the perturbation introduced by RNA regulators, where each input-output interaction depletes one molecule, suggests a simple strategy for insulation: include mechanisms to compensate the depletion of an RNA regulator as it irreversibly binds to its target, for example by linking this binding event to the production of additional copies of RNA regulator. This strategy, hereby called RNA compensation, can be compared to a positive feedback mechanism, because n-copies of a regulator are produced when n-copies are consumed. In this paper, we explore the idea of building RNA compensators for STAR-based circuits using mathematical models and computations. We show that RNA compensation can restore the function of long STAR-based cascades, and eliminate the effects of signal consumption from the steady state behavior of multi-stationary STAR-based circuits. These computational results are broadly relevant to RNA synthetic biology: STARs have been used to create logic gates, cascades and feed-forward loops in combination with many other RNA-based mechanisms, such as small RNA (sRNA) transcriptional repressors, riboregulators, and CRISPR interference (CRISPRi) systems [15, 23, 22, 24]. We anticipate that an effective insulation strategy is necessary to advance towards complex and dynamic RNA circuits. In addition, STARs with RNA compensation have the potential to be implemented as transcriptional connectors bridging traditional protein-based modules, improving the modularity of well-tuned modules. Thus, we expect that the idea of RNA compensation will improve the composability of a variety of novel biomolecular circuits.

## 2 Results

### 2.1 Modeling the stoichiometric operation of STARs

We first establish a model that captures the stoichiometric nature of STAR-based regulation. In our model, species *X* denotes the STAR molecule, which we assume has a zeroth-order production (rate parameter *θ*) and first-order dilution/degradation (rate parameter *ϕ*). In order to capture the kinetic regulatory mechanisms that involves transcription elongation, pausing, and recovery, we define species *Z* that represents a paused transcription elongation complex. This complex includes the DNA template, the RNA polymerase (RNAP) bound to it, and the nascent RNA transcript that presents a binding domain for the STAR molecule (species *X*). When a free STAR molecule *X* binds to the target RNA within the paused elongation complex *Z*, it generates an activated complex *C*: we assume this process happens with association and dissociation rate constants *κ*^+^ and *κ^−^*, where *κ^−^* is not negligible especially in early stages of transcription. If binding of STAR and nascent RNA is successful, then elongation proceeds and complex *C* is converted into the output RNA transcript *Y*. When the stalled elongation complex *C* no longer obstructs the promoter, the binding site for RNAP is recovered and a new elongation complex *Z* forms. We model this process as a first-order reaction that converts *C* into output *Y* and paused elongation complex *Z* with rate parameter *k_cat_*. We assume the RNAP levels to be sufficiently high so that a new RNAP molecule can immediately bind to free promoter. We summarize this description into a set of chemical reactions:

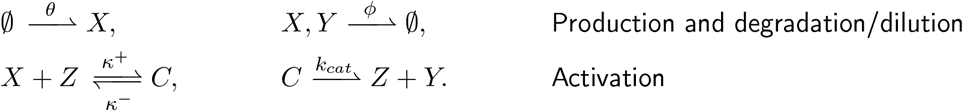

From these reactions we derive an Ordinary Differential Equations (ODE) model by using the law of mass action:

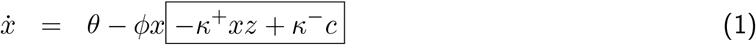

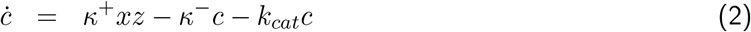

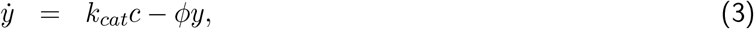

where the sum of paused and activated elongation complexes must be constant, and equal to the total amount of target DNA present: *z* + *c* = *z^tot^*, so that *ż* = −*ċ*.

Equation (1) shows that the level of free STAR molecule *X* is affected by the processes of association and dissociation with the elongation complex, modeled by the terms inside a box. The steady state level of *X* can be found exactly as (detailed derivation in SI section 1.1):

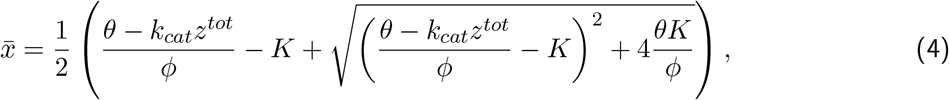

where *K* = (*κ*^−^ + *k_cat_*)/*κ*^+^. In the absence of target binding site (*z^tot^* = 0) the steady state of *X* is 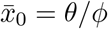. Given a fixed production parameter *θ* (input), the free level of 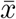 decreases for larger values of *z^tot^*. This effect is due to the fact that the STAR molecule bound to the output RNA is not released after the transcription process is completed, and one STAR molecule is consumed to produce every molecule of output *Y*. The fact that the level of available activator *X* depends on how many downstream binding sites are present is both an opportunity and a challenge: on the one hand, the introduction of binding sites for *X* makes it possible to generate an activation threshold; on the other hand, the presence of uncertain, fluctuating, or increasing binding sites for *X* will perturb the efficiency of the activation pathway. First, we describe how to take advantage of components sequestering *X* to tune the response of the STAR module.

### 2.2 The kinetic and equilibrium response of STAR systems can be adjusted using sequestration and tandem transcription

The tunability of the steady state and kinetic response of a STAR module can be improved by taking advantage of the stoichiometric nature of RNA regulation. First, we can introduce an activation threshold through transcribing an RNA sequester *W* that is complementary to the STAR species *X* (Fig. 1A). Similar sequestration strategies have been demonstrated for STARs and other RNA regulator systems[23]. For simplicity, we assume all RNA species are degraded/diluted with the same reaction parameter *ϕ*. We assume that the double-stranded RNA complex formed by activator *X* hybridized to sequester *W* does not dissociate. Further, the abundance of output *Y* can be tuned by including *n* tandem copies of output *Y*, rather than a single copy. This long transcript could be cleaved into *n* distinct copies of *Y* by leveraging self-cleaving ribozymes[25], or endoribonucleases cleavage sites[26, 27]. When including sequestration and the production of *n*-copies of output the reactions become:

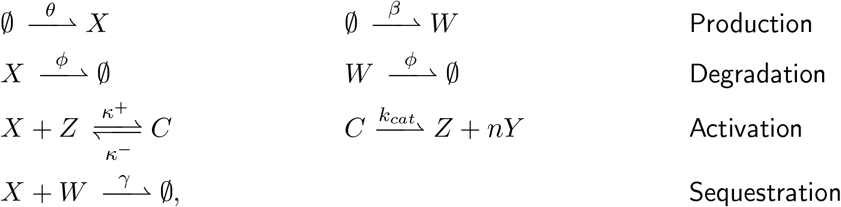

These reactions can be converted to ODEs:

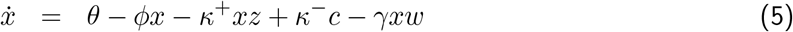

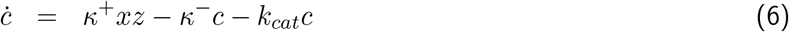

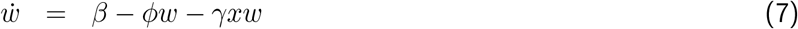

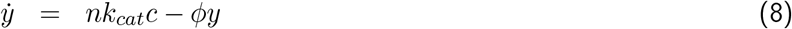

where we assume that the total amount of stalled or active elongation complex remains constant, *z* + *c* = *z^tot^*.

To find the output steady state value of 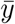 as an exact function of the reaction parameters, one can set equations (5)–(8) equal to zero. This however leads to polynomial equilibrium conditions that are laborious to analyze. To relate the level of *Y* to the production rate constant *θ*, which can be viewed as the input to the reactions, we find expressions relating 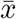 to *θ* and to 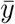, making it possible to generate the map *θ* to 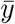 in Fig. 1B by sweeping the values of 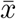 (detailed derivation in SI section 1.2):

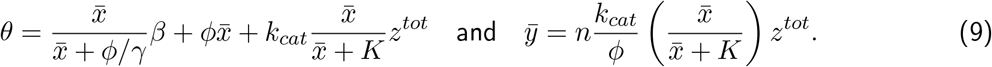

These expressions can be compared with equation (4) expressing 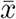 in the absence of sequestration species *W* and tandem transcription of *Y*. If the expected equilibrium level of STAR molecule 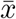 is such that 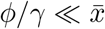 and as a consequence 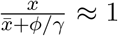, then the level of 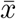 scales with the difference between production rate parameters *θ* and *β*:

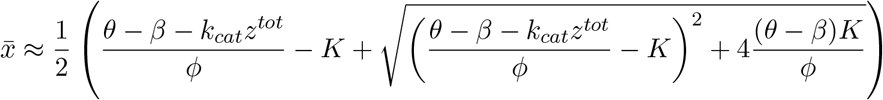

This means that the production parameter *β* of the sequester species serves as a threshold for expression of 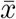 and, in turn, 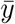. The presence of a threshold is evident in the steady state maps in Fig. 1B. Further, this expression captures the fact that larger amounts of target DNA *z^tot^* cause a reduction of free STAR regulator.

### 2.3 Consumption of STAR molecules can be mitigated using an RNA compensation mechanism

Having created a model that describes the operation and versatility of STAR systems, we turn our attention to the challenges arising when a STAR molecule is used to regulate multiple components downstream and is thereby consumed by multiple processes. We illustrate this by modeling the effect of a single additional target construct, or *load*, *Z_L_* (Fig. 2A and B); for simplicity we assume the STAR species binds and unbinds to the new target with the same reaction rate parameters characterizing the nominal target. This introduces the new chemical reactions:

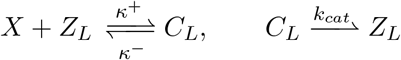

**Figure 2:**
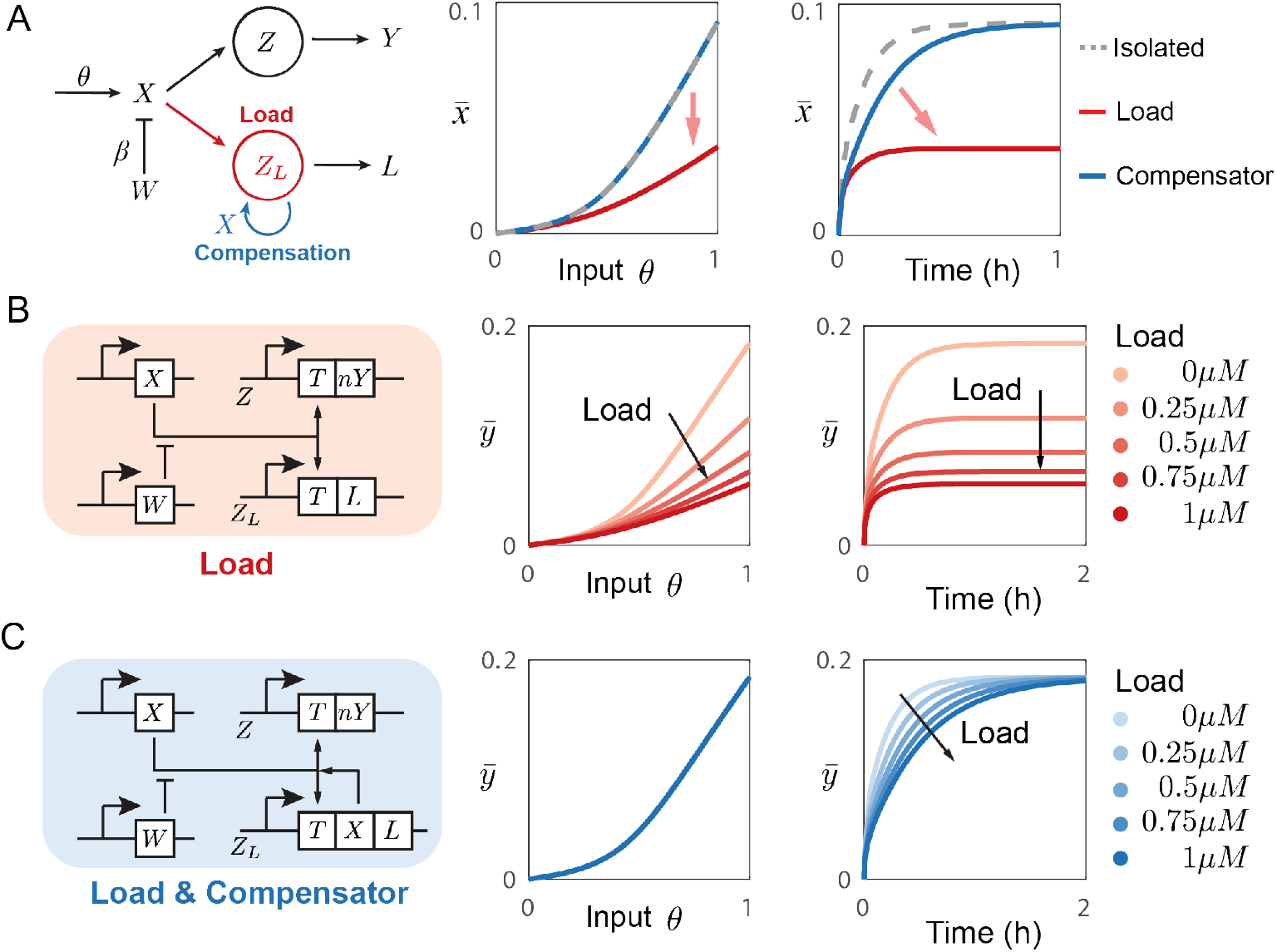
RNA compensation effectively mitigates the consumption of regulator within the STAR system. (A) A single STAR molecule X is consumed in each transcription activation event, leading to a decrease in STAR concentration when additional load is applied. This consumption can be mitigated by replenishing the consumed STAR through a single positive feedback (RNA compensator). (B) When additional load Z_L_ is added to the system, the target output Y is lowered. The extend of reduction depends on the amount of load. (C) The RNA compensator is implemented to the system by adding a STAR following each load. The RNA compensator fully recovers the desired operation of the system in steady state. The delay of response can be observed when additional load Z_L_ is introduced, which functions as extra binding site for X thus more X needs to be transcribed before the system reaches equilibria.

Model (5)–(8) becomes:

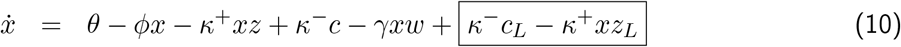

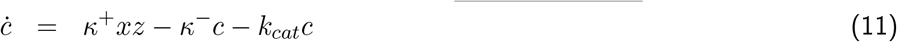

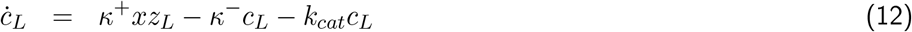

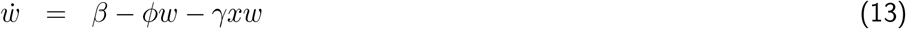

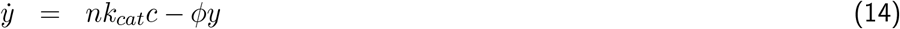

where we assume that the total amount of DNA complex remains constant, *z* + *c* = *z^tot^*, and 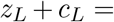 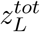. We neglect the output produced by the additional load, and focus on the effects that *Z_L_* has on the primary output species *Y*.

The steady state expressions (9) become:

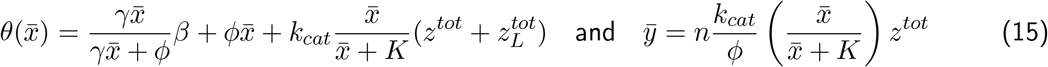

A way to mitigate the depletion of STAR *X* in the presence of additional loads is to supplement STAR production, so that every time a molecule of *X* is bound to the elongating RNA, additional copies of *X* are transcribed. This is equivalent to modifying the activation reaction described earlier:

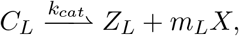

where *m_L_* = 1 considering that an additional copy of *X* is transcribed when the load transcript is generated. This strategy can be implemented through transcriptional fusion of the target RNA, output gene product (*L*) and a STAR (*X*) (Fig. 2A and C), which can be post-transcriptionally separated through the action of self-splicing ribozymes or endoribonucleases that have already been implemented to achieve RNA-only cascade network motifs [15, 19]. When the RNA compensation mechanism is implemented and *m_L_* copies of *X* are added as extra transcription products of activated complex *C*, the dynamics of *X* are modified as:

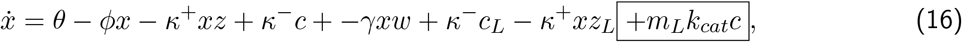

where the compensator term is highlighted in the box. We can find the steady state of compensated *X* (detailed derivation in SI section 1.3),

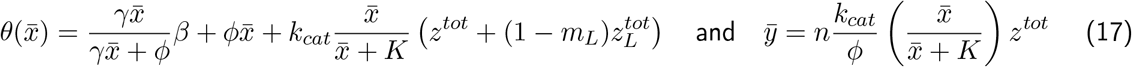

Using this model we simulate the effect of load on the steady state production of STAR (*X*), for varying degrees of load with and without the compensator (Fig. 2B and C, left). We observe a decreasing steady state output of *X* corresponding to the addition of load (Fig. 2B, middle). On the other hand, we observe overlapping outputs of *X* with compensator (Fig. 2C, middle), demonstrating a compensation of the loading effect. Interestingly, the compensation is effective across different levels of load, simulated through increasing the total DNA concentration of binding site *Z_L_*.

We also compare the kinetic response of the uncompensated and compensated systems in the presence of constant input *θ* (Fig. 2B,C right). While the RNA compensator makes it possible to fully recover the desired steady state behavior, we observe a slower response to the presence of input *θ* at time 0. This delay occurs because more *X* is needed for the system to reach the steady state when additional binding site *Z_L_* is introduced.

### 2.4 RNA compensation allows for the creation of long and tunable RNA cascades

Next, we illustrate the usefulness of RNA compensation in simple circuits such as RNA cascades. While cascades of two orthogonal STARs have been created [15], creating longer cascades has been challenging due to signal attenuation. We hypothesize that this attenuation is in part due to RNA consumption at each level of the cascade. Based on this hypothesis, we reason that because our RNA compensation strategy can alleviate signal attenuation, it would allow for the construction of longer RNA cascades.

To test this, we develop a model for an RNA cascade with *q* layers of orthogonal STARs (Fig. 3A). The load within the cascade is contributed by each DNA construct *Z_i_* of the *i^th^* layer. The transcription of *Z_i_* is regulated by the target domain *T_i_*, which consumes a STAR molecule *X_i_* to generate the next-layer STAR molecule *X_i_*_+1_, thereby “passing” the transcriptional signal to the downstream of the cascade. RNA compensation is implemented via an additional domain coding for *X_i_* in tandem with *X_i_*_+1_ within DNA construct *Z_i_* (Fig. 3A). Then we model an external load with the additional DNA construct *Z_Li_*, which consumes STAR *X_i_* and generates output *Y_i_*. Adding *Z_Li_* creates extra loading effect as it competes for STAR *X_i_* with the cascade DNA construct *Z_i_*. Similarly, tandem *X_i_* can be added to *Z_Li_* as compensation mechanism (Fig. 3E). Different load scenarios can be obtained by changing the number *m*, *m_L_*, and 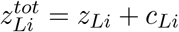 in the following reactions:

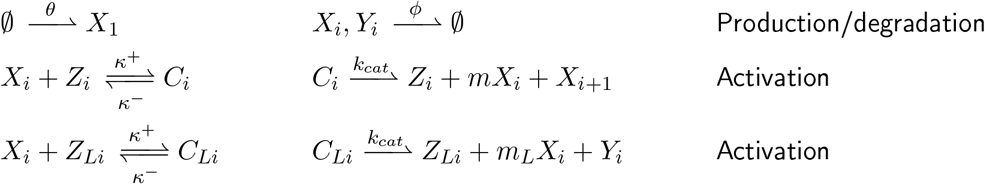

where number *i* annotates the *i^th^* layer of cascade. These reactions can be converted to the ODEs (for 1 < *i* ≤ *q*):

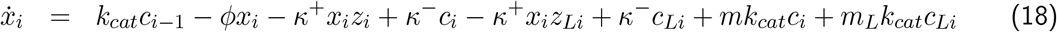

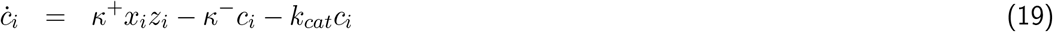

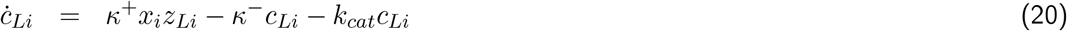

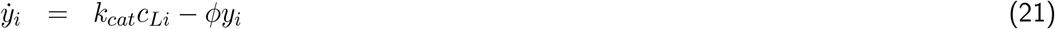

**Figure 3:**
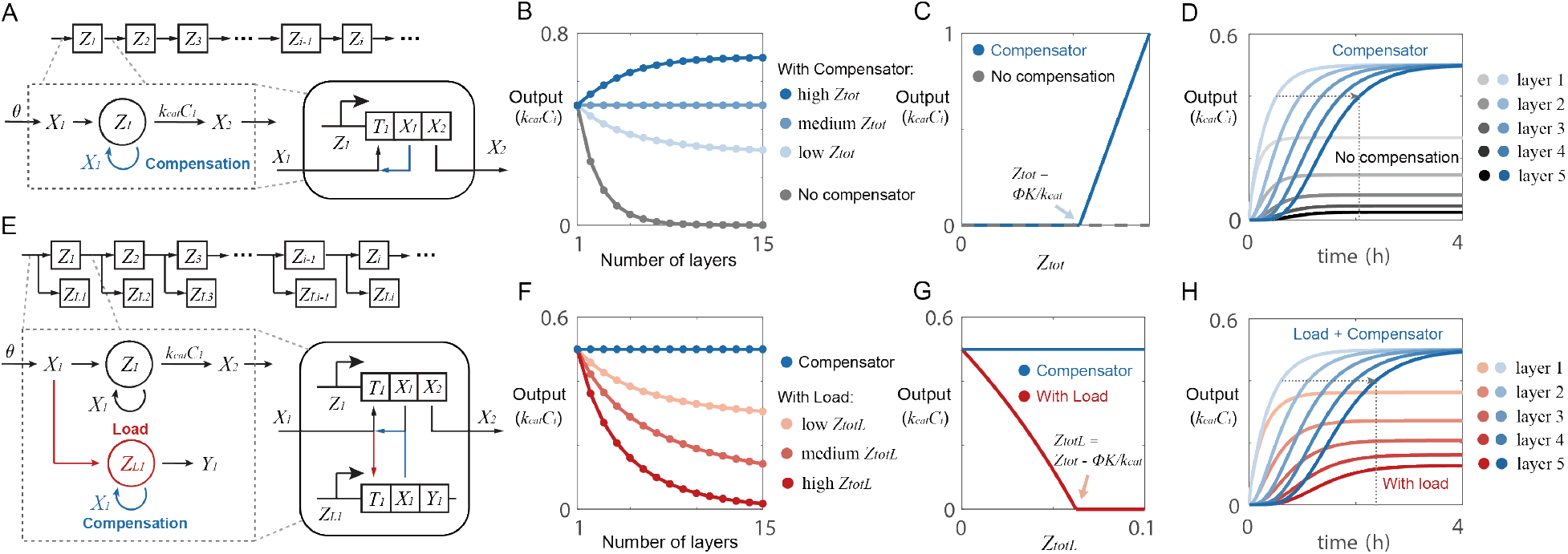
RNA compensator allows for the creation of sustainable and tunable RNA cascades. (A) Schematic of a simple RNA cascade. (B) Simulation of each layer’s transcriptional output k_cat_C_i_ in different cases. When there is no compensation, the signal output of RNA cascade attenuates by each layer and quickly fades out. Adding compensation mechanism stabilizes the output depending on gene concentration z^tot^ (high z^tot^ = 0.7/k_cat_ + ϕK/k_cat_, medium z^tot^ = 0.5/k_cat_ + ϕK/k_cat_, low z^tot^ = 0.3/k_cat_ + ϕK/k_cat_). (C) The prediction of final output when the cascade is long enough to achieve equilibrium output. With compensation, the cascade will have positive stable point if the gene concentration z^tot^ >= ϕK/k_cat_. (D) The dynamic simulation corresponding to the RNA cascade with or without compensation. With no compensation, the output drops quickly after a few layers of the cascade. With compensator, the output can be stabilized. (E) Schematic of an RNA cascade with extra loads applied on each layer. (F) The external load disturbs the cascade through competing for binding of X with cascade nodes (low 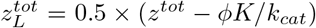, medium 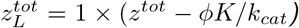, high 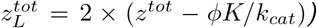. No disturbance is observed in the cascade (with medium level z^tot^) with compensator regardless of the amount of external loads 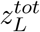. (G) Increasing the concentration of load gene 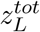 leads to lower equilibrium output of the cascade (become 0 if 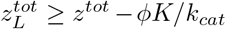) that is mitigated in presence of the compensator. (H) The dynamic simulation corresponding to the RNA cascade with external loads added to each layer. High external loads perturb the ability of the cascade to sustain its nominal output. Adding an RNA compensator to the loads recovers the equilibrium output of the cascade. Meanwhile, delay of response can be observed because the system with additional load needs to accumulate more RNA input to reach equilibrium.

To find the equilibria, we set (18), (19) and (20) equal to 0, obtaining:

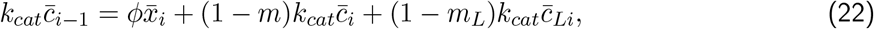

where the initial input *k_cat_c*_0_ = *θ*. Here, we use the transcription rate *k_cat_c_i_* to represent the signal flow of the RNA cascade. When there is no compensation (*m* = 0) nor external loads 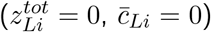 in the cascade, the signal output of *i^th^* layer equals to

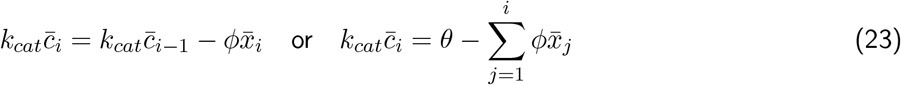

In this case, the maximum of signal flow is limited by the initial input *θ*. The longer the cascade, the lower the output signal will be due to the degradation of *X_i_* in each layer of cascade (Fig. 3B). This signal attenuation effect can be mitigated by setting *m* = 1 (compensation), then the signal outputs becomes (detailed derivation in SI section 2):

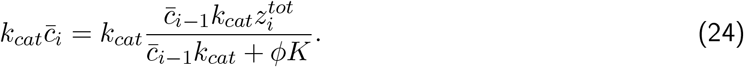

Equation (24) describes a sequence that converges to a finite value when the number of layers *i* is large; this is qualitatively confirmed by the computational simulations in Fig. 3B. If we assume that 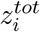 is identical and equal to *z^tot^* for all layers in the cascade, then the value of 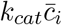 for large *i* must be close to 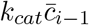. In the limit for *i* → ∞ the equilibrium output of the chain must satisfy *c_i_* = *c*_*i*−1_ in equation (24), and we obtain:

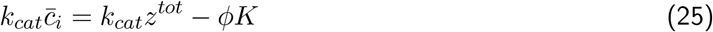

In this case, the output of the system will finally stabilize at *k_cat_z^tot^ − ϕK* (if *z^tot^k_cat_ > ϕK*) or zero (if *z^tot^k_cat_ ≤ ϕK*) (Fig. 3C). Thus, adding a compensation mechanism to the cascade makes it possible for the system to sustain a desired operating level of regulators, which can be tuned by altering the gene concentration *z^tot^*.

Based on the result above, we further investigate the cascade with external loads 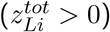 (Fig. 3E). We assume all external loads have same value 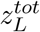 for simplicity. When no compensation is implemented to external loads (*m* = 1, *m_L_* = 0) in equation (22), we can derive the equilibrium output as:

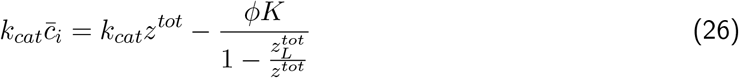

Since 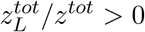 in the presence of external load, the level of 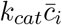 according to equation (26) is lower than the level computed from (25), indicating that the cascade is perturbed by the external load (Fig 3F). When the external load is high 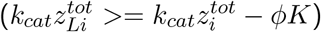, the equilibrium output becomes zero. By implementing the compensation (*m_L_* = 1) in equation (22), the steady state of the system reverts to that computed for the system without external loads, resulting in the full suppression of the effects of load at equilibrium (Fig 3G, SI section 2).

In addition, we investigate how compensation affects the dynamics of the cascade. As expected, in the absence of compensation the output signal quickly attenuates. When the compensator is added, we achieve sustained signal transmission across the RNA cascade (Fig. 3D). When additional load is present in the system, it reduces the steady state output and disrupts the signal transmission. The addition of RNA compensators to the loads makes it possible to recover the nominal steady state output, although (predictably) as we move toward sequential layers in the cascade we observe an increasing delay in achieving the nominal output level (Fig. 3H).

So far, we have shown that the implementation of RNA compensator can significantly improve the composability of STAR cascade, allowing for the construct of more complex RNA circuits. These results prompted us to investigate whether RNA compensation can be used in the context of dynamic circuits to mitigate undesired coupling arising in the presence of loads.

### 2.5 Compensation makes RNA memory elements robust to the presence of downstream loads

We now consider the usefulness of RNA compensation toward the demonstration of bistable RNA-based circuit. Bistable circuits are known to operate as memory elements in many natural processes, and they have been designed with similar goals in engineered biocomputing systems. Although STAR bistable circuits have not yet been demonstrated experimentally, we propose and examine a computational model that relies on self-activation (Grey box, Fig. 4A). In our design, the ON state is achieved through self-activation mediated by a STAR molecule *X*. To compensate for the loss of STAR *X* as it participates in the activation process, STAR repeats are encoded in tandem with the output *Y* in DNA construct *Z*. Importantly, the number of STAR repeats *m* is larger than one in order to achieve self-activation and not simply compensation as discussed above. In order to achieve bistability, we introduce a constitutively expressed sequester molecule *W* to create a tunable activation threshold. We assume the *W* is transcribed and degraded with rate constant *β* and *ϕ*, and binds to *X* with rate constant *γ*.

**Figure 4:**
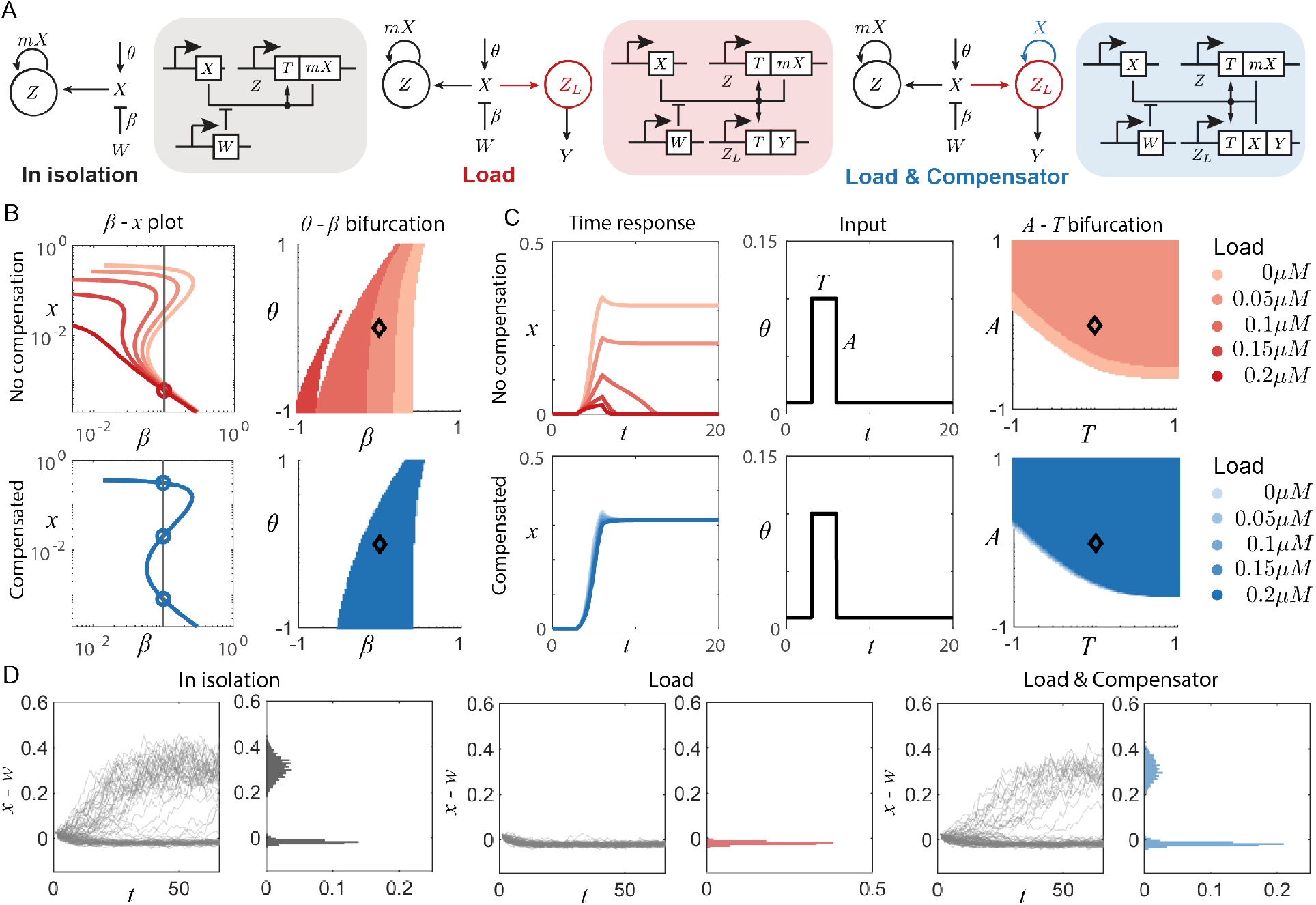
RNA compensator allows for the creation of a robust RNA-based self-activator. A) Schematic and circuit diagram of an RNA self activation memory circuit in isolation (left panel), with load (middle panel), or with load and RNA compensator (right panel). B) Loading analysis of the system in steady state. Left panels are plots of sequester transcription rate (β) versus STAR concentration (x) showing the number of solutions of x at fixed β. Right panels are bifurcation figures of sequester transcription rate (β) and STAR constant transcription rate (θ) showing the shifting of parameter space for bistability with increasing load. The black diamond mark in the center corresponds to the nominal parameter values, which are reported in SI table 1. C) Time-course loading analysis. Left and center panels are time-course response of STAR x by applying an pulse input with amplitude A and time length T. Right panels are A − T specification results to characterize the adequate amplitude A and time length T of input to switch the system from OFF to ON. When the load is higher than 0.1 μM, the systems without compensator (darker reds) are not shown in bifurcation map due to the loss of bistability. D) The RNA compensator effectively mitigates the perturbations in stochastic simulations. Left panels are trajectories of the stochastic simulation where y-axis is the deduction between STAR concentration x and sequester concentration w. Right panels are corresponding histograms at the end of the simulations.

We built a model for this design that can be adapted to consider different load scenarios. We model the presence of downstream loads by including a DNA construct *Z_L_* that encodes the target RNA-controlled domain *T*. This domain introduces load on the self-activation circuit as it depletes STAR molecules *X* (Red box, Fig. 4A). We then model the case in which an RNA compensator is present, and is implemented by encoding an additional copy (*m_L_* = 1) of *X* in tandem with the load output *Y* within the *Z_L_* construct (Blue box, Fig. 4A). These three scenarios can be examined with the single model below:

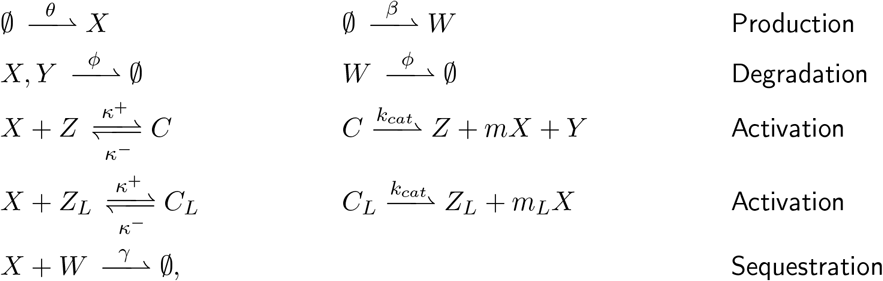

As before, we use law of mass action to derive the ODE model:

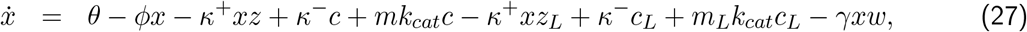

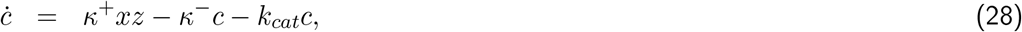

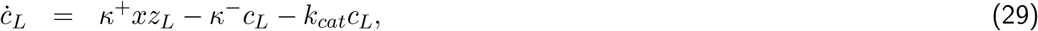

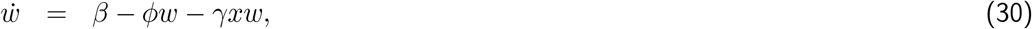

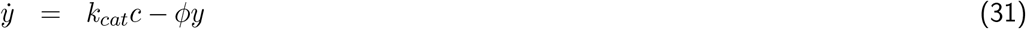

with a mass conservation of the total amount of DNA complex constant, *z* + *c* = *z^tot^*, and 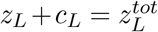

The ability of this model to operate as a memory element can be characterized by seeking the region in parameter space where the circuit exhibits a bistable behavior [28]. Bistability occurs when the system admits three different steady states, two stable and one unstable equilibrium. To check the number of equilibria of system (27)–(30), we found a closed-form expression for the steady state of 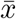 by setting all the above equations equal to zero (see SI Section 3 for details on the derivation).

Based on the method above, we perform steady state analysis on simulations in three different conditions: in the absence of load 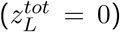, in the presence of load but absence of compensation (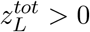, *m_L_* = 0), and in the presence of both load and compensation (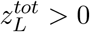, *m_L_* = 1). When the load is present, we examine the behavior of the system at levels of load 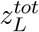 that vary over a four-fold range. To illustrate the results, we plot the the concentration of *X* versus the transcription rate of sequester *β* under different loading conditions, as well as a bifurcation plot showing in color the values of *θ* (transcription rate of *X*) and *β* at which the system is bistable (Fig. 4B). The *x* − *β* plot shows the number of equilibria for a particular value of *β* as the number of intersections between the *x* − *β* curve and the black vertical line. We observe that the presence of load causes the system to transition from bistable to monostable (Fig. 4B, top): while three equilibria are present in absence of load, a single intersection is observed under high load. In contrast, when a compensator is present (*m_L_* = 1), we observe three stable equilibria for any value of load considered 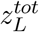 (Fig. 4B, bottom).

While the previous results demonstrate successful load mitigation at steady state, we next sought to understand the effect of the load on the system’s response to time-varying inputs. To do this, we perform numerical simulations to characterize how the self-activation circuit responds to input pulses, introduced via a discrete change of *θ* (transcription rate of *X*) from zero to a maximum value *A* during a finite time interval *T*. Under this transient input, a memory element operating correctly would transition (memory “flip”) from an OFF to an ON equilibrium, remaining committed to the ON equilibrium even when the input is removed. As before, we examine whether the correct operation changes in the presence of load, and whether compensation can mitigate adverse effects of the load. Fig. 4C, top left panel, shows the trajectories of the system’s response to a particular pulse input (top middle) under different loads. We then vary the duration *T* and maximum value *A* of the pulse, showing what input types cause a successful flip of the memory circuit from OFF to ON value (top right). As expected, these results collectively show that when more load is present (top), memory flipping requires an input with a higher peak *A* and longer duration *T*. When the load is higher than 0.1 *μM*, the flipping behavior is not shown in the *A* − *T* bifurcation figure due to the loss of bistability. In contrast (bottom) when RNA compensation is present, the load effects are mitigated, and memory flipping is achieved with inputs having almost identical temporal features regardless of whether the load is present or absent.

Finally, we explore with computations the behavior of the memory element in a stochastic regime using the Gillespie algorithm [29] in Fig. 4D. We consider the same scenarios examined earlier, but we report the difference between the level of *X* and the level of sequester *W* as a measure of the ON/OFF state. We report an ensemble of temporal trajectories, as well as histograms of the stationary levels. In each scenario, initial conditions were set to fall in the unstable region (*x*=0.025, estimated from the deterministic bistability analysis in Fig. 4B). When the load is present, the stationary distribution of the trajectories is shifted toward the OFF state (Fig. 4, middle panels). When RNA compensation is present, the trajectories recover a stationary distribution (right panels) that is similar to the one of the system in isolation (left panels). A slight shift to OFF state is observed in the compensated system compared to the isolated system that suggests a small ratio of compensated systems fails to be turned ON due to the delay generated by the compensation process.

In summary, we observed that RNA compensation makes it possible for the memory circuit to operate robustly in the presence of loads, matching its performance in the absence of load. This suggests that an RNA compensated memory circuit can be used as a module whose specifications are not compromised when interconnected to other devices.

## 3 Discussion

We describe a strategy to improve the composability of RNA transcriptional circuits based upon positive feedback. Our strategy, which we termed RNA compensation, addresses the stoichiometric consumption of RNA regulator molecules that occurs within the small transcription activating RNA (STAR) regulatory operation. RNA compensators mitigate input loss by production of additional RNA regulatory inputs in tandem with each output gene, thus replenishing the RNA input pool. We demonstrated through computations and mathematical analysis that RNA compensators can fully recover the steady state response of STAR regulators when connected to multiple downstream modules, and greatly increase their modularity as synthetic biology parts. RNA compensators also improve the connectivity and scalability of RNA signaling cascades, as well as the robustness and tunability of a model for bistable RNA self-activation.

Our RNA compensation strategy provides a novel and distinct insulation approach for addressing signal consumption in genetic circuits. To date, research has largely focused on the problem of reducing a particular form of perturbation known as retroactivity, using strategies based on negative feedback [6]. In brief, these systems harness a high-gain negative feedback from the output. As the gain grows, the output become close to the ratio between the input and feedback strength, which is independent of retroactivity [4]. Theoretical and experimental analysis have demonstrated negative feedback to be a robust strategy to address retroactivity [9]. However, one challenge with this strategy is that it requires high-level expression of an inhibitor molecule, which increase circuit complexity and metabolic burden. Counter to this, a major potential advantage of RNA compensation is its simplicity, requiring transcriptional fusion of an RNA input to each output gene. As a result, this strategy is genetically compact, requiring only ~ 100 nt of additional sequence to be added for each output. Additionally, it is metabolically efficient, requiring only the minimum number of molecules to achieve compensation. Of course, an inherent limitation of this strategy is the requirement of a regulator system with a gain equal to 1, meaning for each output an input is consumed. While this means the strategy is not readily applicable to all regulatory mechanisms, for example, protein-based transcription factors, it is ideally suited to address retroactivity for RNA regulators that rely upon formation of extended duplexes that are largely irreversible.

The RNA compensator represents a synthetic approach to achieve concentration robustness against the presence of output loads. It shares similarities to absolute concentration robustness (ACR) systems. ACR occurs when a biological system can maintain a molecular species at fixed concentration regardless of the initial conditions [30]. In nature, several molecular control systems have been described that can achieve ACR, for example, the EnvZ/OmpR two-component system of *E. coli* [31]. In this system, EnvZ serves to phosphorylate and desphorylate the response regulator OmpR. Under cellular conditions, the phosphorated OmpR (OmpR-P) is insensitive to both variations in OmpR and EnvZ, meaning ACR of OmpR-P is achieved [32]. This is implemented catalytically through the bifunctional enzymes acting as both kinase and phosphatase. Other examples of ACR found in nature include the glyoxylate bypass regulation system that is also implemented through the action of catalytic biomolecules [33]. Like ACR systems, the RNA compensator leads to fixed STAR concentrations that depend solely on its production and degradation rate, thus forming a concentration robustness towards downstream loads. The RNA compensator represents a novel and synthetic implementation of concentration robustness that acts at the level of gene expression.

We anticipate that the experimental implementation of RNA compensators will allow for further advances in RNA circuitry, particularly multi-node RNA cascades and dynamic RNA-based systems similar to those created using protein-based regulators [34]. As dynamic circuits are sensitive to disturbances and perturbations, any downstream module, even a single reporter, can drastically alter the behavior of the dynamic circuit [35, 5]. By implementing RNA compensators as connectors between modules, perturbations introduced by downstream modules can be greatly mitigated, allowing for the assembly of well-tuned dynamic circuits without additional tuning.

In summary, we describe a novel RNA-based approach to mitigate retroactivity. Our compensator approach adds the available strategies to mitigate retroactivity within biological systems, which we anticipate will be of value for RNA-based synthetic systems.

## Acknowledgements

EF and CCS were partially supported by NSF-BBSRC/BIO award 2020039. JC and BL were partially supported by Alfred P. Sloan Reasearch Fellowship [FG-2018-10500] and NSF award 2124306.

## References

[1] Gardner TS, Cantor CR, Collins JJ. Construction of a genetic toggle switch in Escherichia coli. Nature. 2000 1;403:339–342.

[2] Santos-Moreno J, Tasiudi E, Stelling J, Schaerli Y. Multistable and dynamic CRISPRi-based synthetic circuits. Nature Communications. 2020;11.

[3] Saez-Rodriguez J, Kremling A, Gilles ED. Dissecting the puzzle of life: modularization of signal transduction networks. Computers & chemical engineering. 2005;29(3):619–629.

[4] Vecchio DD, Ninfa AJ, Sontag ED. Modular cell biology: Retroactivity and insulation. Molecular Systems Biology. 2008;4.

[5] Franco E, Friedrichs E, Kim J, Jungmann R, Murray R, Winfree E, et al. Timing molecular motion and production with a synthetic transcriptional clock. Proceedings of the National Academy of Sciences of the United States of America. 2011;108.

[6] McBride C, Shah R, Vecchio DD. The Effect of Loads in Molecular Communications. Proceedings of the IEEE. 2019;107.

[7] Vecchio DD. Modularity, context-dependence, and insulation in engineered biological circuits; 2015.

[8] Nilgiriwala KS, Jiménez J, Rivera PM, Vecchio DD. Synthetic Tunable Amplifying Buffer Circuit in E. coli. ACS Synthetic Biology. 2015;4.

[9] Huang HH, Bellato M, Qian Y, Cárdenas P, Pasotti L, Magni P, et al. dCas9 regulator to neutralize competition in CRISPRi circuits. Nature Communications. 2021;12.

[10] Mishra D, Rivera PM, Lin A, Vecchio DD, Weiss R. A load driver device for engineering modularity in biological networks. Nature Biotechnology. 2014;32.

[11] Aoki SK, Lillacci G, Gupta A, Baumschlager A, Schweingruber D, Khammash M. A universal biomolecular integral feedback controller for robust perfect adaptation. Nature. 2019;570.

[12] Huang HH, Qian Y, Vecchio DD. A quasi-integral controller for adaptation of genetic modules to variable ribosome demand. Nature Communications. 2018;9.

[13] Vecchio DD, Dy AJ, Qian Y. Control theory meets synthetic biology; 2016.

[14] Chappell J, Watters KE, Takahashi MK, Lucks JB. A renaissance in RNA synthetic biology: New mechanisms, applications and tools for the future; 2015.

[15] Chappell J, Westbrook A, Verosloff M, Lucks JB. Computational design of small transcription activating RNAs for versatile and dynamic gene regulation. Nature Communications. 2017 12;8.

[16] Gander MW, Vrana JD, Voje WE, Carothers JM, Klavins E. Digital logic circuits in yeast with CRISPR-dCas9 NOR gates. Nature Communications. 2017 5;8.

[17] Green AA, Kim J, Ma D, Silver PA, Collins JJ, Yin P. Complex cellular logic computation using ribocomputing devices. Nature. 2017 8;548:117–121.

[18] Kuo J, Yuan R, Sánchez C, Paulsson J, Silver PA. Toward a translationally independent RNA-based synthetic oscillator using deactivated CRISPR-Cas. Nucleic Acids Research. 2020;48.

[19] Lucks JB, Qi L, Mutalik VK, Wang D, Arkin AP. Versatile RNA-sensing transcriptional regulators for engineering genetic networks. Proceedings of the National Academy of Sciences of the United States of America. 2011;108.

[20] Takahashi MK, Chappell J, Hayes CA, Sun ZZ, Kim J, Singhal V, et al. Rapidly Characterizing the Fast Dynamics of RNA Genetic Circuitry with Cell-Free Transcription-Translation (TX-TL) Systems. ACS Synthetic Biology. 2015;4.

[21] Hu CY, Takahashi MK, Zhang Y, Lucks JB. Engineering a Functional Small RNA Negative Autoregulation Network with Model-Guided Design. ACS Synthetic Biology. 2018;7.

[22] Westbrook A, Tang X, Marshall R, Maxwell CS, Chappell J, Agrawal DK, et al. Distinct timescales of RNA regulators enable the construction of a genetic pulse generator. Biotechnology and Bioengineering. 2019;116.

[23] Lee YJ, Kim SJ, Moon TS. Multilevel Regulation of Bacterial Gene Expression with the Combined STAR and Antisense RNA System. ACS Synthetic Biology. 2018;7.

[24] Greco FV, Pandi A, Erb TJ, Grierson CS, Gorochowski TE. Harnessing the central dogma for stringent multi-level control of gene expression. Nature Communications. 2021;12.

[25] Lou C, Stanton B, Chen YJ, Munsky B, Voigt CA. Ribozyme-based insulator parts buffer synthetic circuits from genetic context. Nature Biotechnology. 2012;30.

[26] Haurwitz RE, Jinek M, Wiedenheft B, Zhou K, Doudna JA. Sequence- and structure-specific RNA processing by a CRISPR endonuclease. Science. 2010;329.

[27] Ferreira R, Skrekas C, Nielsen J, David F. Multiplexed CRISPR/Cas9 Genome Editing and Gene Regulation Using Csy4 in Saccharomyces cerevisiae. ACS Synthetic Biology. 2018;7.

[28] Inniss MC, Silver PA. Building synthetic memory; 2013.

[29] Gillespie DT. Exact stochastic simulation of coupled chemical reactions. vol. 81; 1977. .

[30] Shinar G, Feinberg M. Structural sources of robustness in biochemical reaction networks. Science. 2010;327.

[31] Batchelor E, Goulian M. Robustness and the cycle of phosphorylation and dephosphorylation in a two-component regulatory system. Proceedings of the National Academy of Sciences of the United States of America. 2003;100.

[32] Cappelletti D, Gupta A, Khammash M. A hidden integral structure endows absolute concentration robust systems with resilience to dynamical concentration disturbances. Journal of the Royal Society Interface. 2020;17.

[33] LaPorte DC, Thornsness PE, Koshland DE. Compensatory phosphorylation of isocitrate dehydrogenase. A mechanism for adaptation to the intracellular environment. Journal of Biological Chemistry. 1985;260.

[34] Elowitz MB, Leibier S. A synthetic oscillatory network of transcriptional regulators. Nature. 2000 1;403:335–338.

[35] Jayanthi S, Nilgiriwala KS, Vecchio DD. Retroactivity controls the temporal dynamics of gene transcription. ACS Synthetic Biology. 2013;2.

