## Supplementary material for "RNA compensation: A positive feedback insulation strategy for RNA-based networks": SI

### Supplementary Information - RNA compensator: A positive feedback insulation strategy for RNA-based networks

October 26, 2021

**2 Derivation of equilibrium points in RNA cascades** **5**

**3 Modeling of an RNA-based memory system** **7**

**4 Table 1 - General parameters** **8**

### 1 STAR module analysis

#### 1.1 Model derivation of the small transcription activating RNAs (STAR) system

In this section, we perform a steady state analysis of the STAR system. To help the reader, we summarize the chemical reaction,

$$\emptyset \xrightarrow{\theta} X \qquad X, Y \xrightarrow{\phi} \emptyset \qquad \text{Production}$$

$$X + Z \underset{\kappa^-}{\overset{\kappa^+}{\rightleftharpoons}} C \qquad C \xrightarrow{k_{cat}} Z + nY. \qquad \text{Activation}$$

We derive the Ordinary Differential Equations (ODEs) by using the law of mass action:

$$\dot{x} = \theta - \phi x - \kappa^+ xz + \kappa^- c \quad (1)$$

$$\dot{c} = \kappa^+ xz - \kappa^- c - k_{cat} c \quad (2)$$

$$\dot{y} = nk_{cat} c - \phi y \quad (3)$$

with a mass conservation of the total amount of DNA complex constant, $z + c = z^{tot}$.

**Steady state** To find the steady state, we make equations (1)-(3) equal to zero. From $\dot{c} = 0$, we obtain

$$\bar{c} = \frac{\bar{x}}{(\kappa^- + k_{cat})/\kappa^+}\bar{z} = \frac{\bar{x}}{K}\bar{z}, \quad (4)$$

where we define the dissociation constant $K = (\kappa^- + k_{cat})/\kappa^+$. Next, we use mass conservation, $\bar{z} + \bar{c} = z^{tot}$, and find that

$$\bar{c} = \frac{\bar{x}}{\bar{x} + K} z^{tot}. \quad (5)$$

From $\dot{x} = 0$, and $\dot{y} = 0$, we plug $\bar{c}$ in these equations, and leads to

$$0 = \theta - \phi\bar{x} - k_{cat} z^{tot} \frac{\bar{x}}{\bar{x} + K} \quad (6)$$

$$0 = nk_{cat} z^{tot} \frac{\bar{x}}{\bar{x} + K} - \phi\bar{y} \quad (7)$$

Equation (6) leads to a second order polynomial, $\bar{x}^2 + p_1\bar{x} - p_0 = 0$, where $p_1 = K + (k_{cat}z^{tot} - \theta)/\phi$ and $p_0 = \theta K/\phi$. This result in a single positive real solution and we can find

$$\bar{x} = \frac{-p_1 + \sqrt{p_1^2 + 4p_0}}{2} \quad \text{and} \quad \bar{y} = n\frac{k_{cat} z^{tot}}{\phi}\left(\frac{\bar{x}}{\bar{x} + K}\right). \quad (8)$$

**Stability analysis** We compute the Jacobian matrix for the sytem (1)-(3), resulting in

$$J = \begin{bmatrix} -\phi - \kappa^+\bar{z} & \kappa^+\bar{x} + \kappa^- & 0 \\ \kappa^+\bar{z} & -\kappa^+\bar{x} - \kappa^- - k_{cat} & 0 \\ 0 & k_{cat} & -\phi \end{bmatrix}. \quad (9)$$

The system can only admit negative eigen values when the following condition is satisfied,

$$\phi k^+\bar{x} + \phi k^- + \phi k_{cat} > -k_{cat}k^+\bar{z}.$$

Since this condition is satisfied by all parameters, we conclude that the system is unconditional stable.

#### 1.2 Molecular sequestration tunes the dose response of the STAR system

From the description of main text, we summarize the chemical reaction,

$$\begin{array}{lll} \emptyset \xrightarrow{\theta} X & X, Y \xrightarrow{\phi} \emptyset & \text{Production/Degradation} \\ \emptyset \xrightarrow{\beta} W & W \xrightarrow{\phi} \emptyset & \text{Production/Degradation} \\ X + Z \underset{\kappa^-}{\overset{\kappa^+}{\rightleftharpoons}} C & C \xrightarrow{k_{cat}} Z + nY & \text{Activation} \\ X + W \xrightarrow{\gamma} \emptyset. & & \text{Sequestration} \end{array}$$

We derive the Ordinary Differential Equations (ODEs) by using the law of mass action:

$$\dot{x} = \theta - \phi x - \kappa^+ xz + \kappa^- c - \gamma xw \quad (10)$$

$$\dot{c} = \kappa^+ xz - \kappa^- c - k_{cat} c \quad (11)$$

$$\dot{w} = \beta - \gamma xw - \phi w \quad (12)$$

$$\dot{y} = nk_{cat} c - \phi y, \quad (13)$$

with a mass conservation of the total amount of DNA complex constant, $z + c = z^{tot}$.

**Steady state** We follow similar steps as before, and by making $\dot{c} = 0$, we find that $\bar{c} = \frac{\bar{x}}{\bar{x}+K} z^{tot}$. Further, by plugging in the other equations $\dot{x} = \dot{w} = \dot{y} = 0$, this results in

$$0 = \theta - \phi\bar{x} - k_{cat} z^{tot} \frac{\bar{x}}{\bar{x} + K} - \gamma\bar{x}\bar{w} \quad (14)$$

$$0 = \beta - \phi\bar{w} - \gamma\bar{x}\bar{w} \quad (15)$$

$$0 = nk_{cat} z^{tot} \frac{\bar{x}}{\bar{x} + K} - \phi\bar{y}. \quad (16)$$

From equation (14)-(15), we can find $\bar{w}$ as a funtion of $\bar{x}$

$$\bar{w} = \frac{\theta - \phi\bar{x} - k_{cat} z^{tot} \frac{\bar{x}}{\bar{x}+K}}{\gamma\bar{x}} = \frac{\beta}{\gamma\bar{x} + \phi}. \quad (17)$$

This leads to a third order polynomial. However, a simple way to find the input-output is to find $\theta$ as a function of $\bar{x}$. This results in

$$\theta(\bar{x}) = \frac{\gamma\bar{x}}{\gamma\bar{x} + \phi}\beta + \phi\bar{x} + k_{cat} z^{tot} \frac{\bar{x}}{\bar{x} + K}. \quad (18)$$

**Stability analysis.** We compute the Jacobian matrix for the system (10)-(13), resulting in

$$J = \begin{bmatrix} -\phi - \kappa^+\bar{z} - \gamma\bar{w} & \kappa^+\bar{x} + \kappa^- & -\gamma\bar{x} & 0 \\ \kappa^+\bar{z} & -\kappa^+\bar{x} - \kappa^- - k_{cat} & 0 & 0 \\ -\gamma\bar{w} & 0 & -\phi - \gamma\bar{x} & 0 \\ 0 & 0 & nk_{cat} & -\phi \end{bmatrix}. \quad (19)$$

The system $J$ can admit negative eigenvalues if the following conditions are satisfied,

$$(2\phi + \gamma(\bar{w} + \bar{x}))(\kappa^+\bar{x} + \kappa^- + k_{cat}) + (\phi + \kappa^+\bar{z} + \gamma\bar{w})\phi + (\phi + \kappa^+\bar{z})(\gamma\bar{x}) > -k_{cat}\kappa^+\bar{z}$$

and

$$\phi(\phi + \gamma(\bar{x} + \bar{w}))(\kappa^+\bar{x} + \kappa^- + k_{cat}) > -k_{cat}\kappa^+\bar{z}(\phi + \gamma\bar{x}).$$

Then, we can conclude that the STAR system that incorporates sequestration is unconditional stable for any parameter value.

#### 1.3 RNA compensator mitigates the loading effects

In section we will consider the activation of two downstream process. Now the activator can bind and unbind to $Z$ and $Z_L$, forming complex $C$ and $C_L$. $C$ produces a copy of $X$ and $Y$, which are separated by insulators such as ribozymes or endoribonuclease sites. Meanwhile, $C_L$ produces $m_L$ copies of $X$ using same insulation strategy. We set $m_L = 0$ to simulate loading effect and set $m_L = 1$ when RNA compensator is implemented. Both $X$ and $Y$ degrade at a constant rate $\phi$. We summarize the chemical reaction as

$$
\begin{array}{lll}
\emptyset \xrightarrow{\theta} X & X, Y \xrightarrow{\phi} \emptyset & \text{Production/Degradation} \\
\emptyset \xrightarrow{\beta} W & W \xrightarrow{\phi} \emptyset & \text{Production/Degradation} \\
X + Z \underset{\kappa^-}{\overset{\kappa^+}{\rightleftharpoons}} C & C \xrightarrow{k_{cat}} Z + X + nY. & \text{Activation} \\
X + Z_L \underset{\kappa^-}{\overset{\kappa^+}{\rightleftharpoons}} C_L & C_L \xrightarrow{k_{cat}} Z_L + m_L X. & \text{Activation} \\
X + W \xrightarrow{\gamma} \emptyset. & & \text{Sequestration}
\end{array}
$$

We derive the Ordinary Differential Equations (ODEs) by using the law of mass action:

$$
\begin{aligned}
\dot{x} &= \theta - \phi x - \gamma x w - \kappa^+ x z + \kappa^- c - \kappa^+ x z_L + \kappa^- c_L + m_L k_{cat} c_L & (20) \\
\dot{c} &= \kappa^+ x z - \kappa^- c - k_{cat} c & (21) \\
\dot{c}_L &= \kappa^+ x z_L - \kappa^- c_L - k_{cat} c_L & (22) \\
\dot{w} &= \beta - \phi w - \gamma x w & (23) \\
\dot{y} &= n k_{cat} c - \phi y & (24)
\end{aligned}
$$

with a mass conservation of the total amount of DNA complex constant, $z + c = z^{tot}$ and $z_L + c_L = z_L^{tot}$.

**Steady state analysis.** By making $\dot{c} = \dot{c}_L = 0$, following similar steps as before, we find that $\bar{c} = x z^{tot}/(x + K)$, and $\bar{c}_L = x z_L^{tot}/(x + K)$,

$$
\begin{aligned}
0 &= \theta - \phi \bar{x} - \gamma \bar{x} \bar{w} - k_{cat} z^{tot} \frac{\bar{x}}{\bar{x} + K} + (m_L - 1) k_{cat} z_L^{tot} \frac{\bar{x}}{\bar{x} + K} & (25) \\
0 &= \beta - \phi \bar{w} - \gamma \bar{x} \bar{w} & (26) \\
0 &= n k_{cat} z^{tot} \frac{\bar{x}}{\bar{x} + K} - \phi y & (27)
\end{aligned}
$$

From equation (25)-(26), we can find $\bar{w}$ as a funtion of $\bar{x}$

$$
\bar{w} = \frac{\theta - \phi \bar{x} - k_{cat} z^{tot} \frac{\bar{x}}{\bar{x}+K} + (m_L - 1) k_{cat} z_L^{tot} \frac{\bar{x}}{\bar{x}+K}}{\gamma \bar{x}} = \frac{\beta}{\gamma \bar{x} + \phi}. \tag{28}
$$

This leads to a third order polynomial. However, a simple way to find the input-output is to find $\theta$ as a function of $\bar{x}$. This results in

$$
\theta(\bar{x}) = \frac{\gamma \bar{x}}{\gamma \bar{x} + \phi} \beta + \phi \bar{x} + \left\{ z^{tot} + (1 - m_L) z_L^{tot} \right\} k_{cat} \frac{\bar{x}}{\bar{x} + K}, \tag{29}
$$

and $\bar{y} = n k_{cat} z^{tot} \frac{\bar{x}}{\bar{x}+K}$.

**Stability analysis.** We write down the Jacobian matrix for the system (20)-(24), resulting in

$$J = \begin{bmatrix} -\phi - \kappa^+(\bar{z} + \bar{z}_L) - \gamma\bar{w} & \kappa^- + k^+\bar{x} & \kappa^- + m_L k_{cat} + k^+\bar{x} & -\gamma\bar{x} & 0 \\ \kappa^+\bar{z} & -\kappa^- - k^+\bar{x} & 0 & 0 & 0 \\ \kappa^+\bar{z}_L & 0 & -\kappa^- - m_L k_{cat} - k^+\bar{x} & 0 & 0 \\ -\gamma\bar{w} & 0 & 0 & -\phi - \gamma\bar{x} & \\ 0 & nk_{cat} & 0 & 0 & \phi \end{bmatrix}. \tag{30}$$

We evaluate the stability of the system by computing the eigenvalues of J.

### 2 Derivation of equilibrium points in RNA cascades

A multi-layer RNA cascade is described as following reactions:

| | | |
|---|---|---|
| $\emptyset \xrightarrow{\theta} X_1$ | Initial input | |
| $X_i \xrightarrow{\phi} \emptyset$ | $Y_i \xrightarrow{\phi} \emptyset$ | Degradation |
| $X_i + Z_i \underset{\kappa^-}{\overset{\kappa^+}{\rightleftharpoons}} C_i$ | $C_i \xrightarrow{k_{cat}} Z_i + mX_i + X_{i+1}$ | Activation |
| $X_i + Z_{Li} \underset{\kappa^-}{\overset{\kappa^+}{\rightleftharpoons}} C_{Li}$ | $C_{Li} \xrightarrow{k_{cat}} Z_{Li} + m_L X_i + Y_i$ | Load |

These reactions are converted to ODEs:

$$\dot{x}_i = k_{cat}c_{i-1} - \phi x_i - \kappa^+ x_i z_i + \kappa^- c_i - \kappa^+ x_i z_{Li} + \kappa^- c_{Li} + mk_{cat}c_i + m_L k_{cat} c_{Li} \tag{31}$$

$$\dot{c}_i = \kappa^+ x_i z_i - \kappa^- c_i - k_{cat} c_i \tag{32}$$

$$\dot{c}_{Li} = \kappa^+ x_i z_{Li} - \kappa^- c_{Li} - k_{cat} c_{Li} \tag{33}$$

$$\dot{y}_i = k_{cat} c_{Li} - \phi y_i \tag{34}$$

where we define $z_i^{tot} = c_i + z_i$, $z_{Li}^{tot} = c_L i + z_{Li}$, and $K = \frac{\kappa^- + k_{cat}}{\kappa^+}$. By setting (32) equal to 0 we derive:

$$\bar{c}_i = \frac{\bar{x}_i \bar{z}_i^{tot}}{\bar{x}_i + K} \quad \text{as well as} \quad \bar{x}_i = \frac{K\bar{c}_i}{\bar{z}_i^{tot} - \bar{c}_i} \tag{35}$$

Setting (31) equal to 0, we get

$$k_{cat}\bar{c}_{i-1} = \phi\bar{x}_i + (1-m)k_{cat}\bar{c}_i + (1-m_L)k_{cat}\bar{c}_{Li} \tag{36}$$

When there is no compensation ($m = 0$) nor additional loads ($z_{Li}^{tot} = 0$, $\bar{c}_{Li} = 0$) in the cascade, the relation between the transcription output of $i^{th}$ layer and $i - 1^{th}$ layer is

$$k_{cat}\bar{c}_i = k_{cat}\bar{c}_{i-1} - \phi\bar{x}_i \tag{37}$$

Consider that the initial input $k_{cat}c_0 = \theta$, the equation above can be written as

$$k_{cat}\bar{c}_i = \theta - \sum_{n=1}^{i} \phi\bar{x}_n \tag{38}$$

This suggests that in an RNA cascade without compensation, the maximum of signal output is limited by the initial input $\theta$. The longer the cascade is, the lower the output signal will be due to the depletion of $X_i$ in each layer of cascade.

Next, we solve $k_{cat}\bar{c}_i$ with (35) and (37), and we get:

$$k_{cat}\bar{c}_i = k_{cat}\bar{c}_{i-1} - \frac{\phi K \bar{c}_i}{z_i^{tot} - \bar{c}_i} \tag{39}$$

$$k_{cat}\bar{c}_i = \frac{p_1 - \sqrt{p_1^2 - 4k_{cat}^2\bar{c}_{i-1}z_i^{tot}}}{2} \quad \text{where} \quad p_1 = k_{cat}\bar{c}_{i-1} + k_{cat}z_i^{tot} + \phi K \tag{40}$$

Then we add the compensator by setting $m = 1$ in (36) and replacing $x_i$ with (35). Then, the transcription output becomes:

$$k_{cat}\bar{c}_i = k_{cat}\frac{\bar{c}_{i-1}k_{cat}z_i^{tot}}{\bar{c}_{i-1}k_{cat} + \phi K} \tag{41}$$

This is a convergent sequence because $(\bar{c}_{i-1}k_{cat})/(\bar{c}_{i-1}k_{cat} + \phi K) < 1$ and $k_{cat}\bar{c}_i < k_{cat}z_i^{tot}$. So we can find the limit by letting the $k_{cat}c_i = k_{cat}c_{i-1}$, and we get the equilibrium output for a long cascade

$$k_{cat}\bar{c}_i = k_{cat}z_i^{tot} - \phi K \tag{42}$$

Supposing that $z_i^{tot}$ is a fixed number $z^{tot}$ for all layers and the cascade is long enough, the output of the system will finally stabilize at $k_{cat}z^{tot} - \phi K$ (if $z^{tot}k_{cat} > \phi K$) or zero (if $z^{tot}k_{cat} <= \phi K$).

Then we further investigate the loading effect on cascade with external loads ($z_{Li}^{tot} > 0$). When no compensation is implemented ($m_L = 0$) in (36), signal output of $i^{th}$ layer become

$$k_{cat}\bar{c}_i = \frac{p_1 - \sqrt{p_1^2 - 4k_{cat}^2\bar{c}_{i-1}z_{Li}^{tot}}}{2\frac{z_{Li}^{tot}}{z_i^{tot}}} \quad \text{where} \quad p_1 = k_{cat}\bar{c}_{i-1} + k_{cat}\bar{z}_{Li}^{tot} + \phi K \tag{43}$$

Similarly, we get the equilibrium point

$$k_{cat}\bar{c}_i = k_{cat}z_i^{tot} - \frac{\phi K}{1 - \frac{z_{Li}^{tot}}{z_i^{tot}}} \tag{44}$$

The value of $k_{cat}\bar{c}_i$ in (44) is lower than the value computed from (42), indicating that the cascade is disturbed by the external load. When the external load is high ($k_{cat}z_{Li}^{tot} >= k_{cat}z_i^{tot} - \phi K$), the equilibrium point becomes zero. By implementing the compensation ($n = 1$), the system is recovered to (41) and (42), result in fully mitigation of loading effect in equilibrium.

#### 3 Modeling of an RNA-based memory system

To help the readers with the description of the RNA-based bistable design, we summarize the chemical reactions

| | | |
|---|---|---|
| $\emptyset \xrightarrow{\theta} X$ | $X, Y \xrightarrow{\phi} \emptyset$ | Production/Degradation |
| $\emptyset \xrightarrow{\beta} W$ | $W \xrightarrow{\phi} \emptyset$ | Production/Degradation |
| $X + Z \underset{\kappa^-}{\overset{\kappa^+}{\rightleftharpoons}} C$ | $C \xrightarrow{k_{cat}} Z + mX + Y.$ | Activation |
| $X + Z_L \underset{\kappa^-}{\overset{\kappa^+}{\rightleftharpoons}} C_L$ | $C_L \xrightarrow{k_{cat}} Z_L + m_L X.$ | Activation |
| $X + W \xrightarrow{\gamma} \emptyset.$ | | Sequestration |

We derive the Ordinary Differential Equations (ODEs) by using the law of mass action:

$$\dot{x} = \theta - \phi x - \gamma x w - \kappa^+ x z + \kappa^- c + m k_{cat} c - \kappa^+ x z_L + \kappa^- c_L + m_L k_{cat} c_L \quad (45)$$
$$\dot{c} = \kappa^+ x z - \kappa^- c - k_{cat} c \quad (46)$$
$$\dot{c}_L = \kappa^+ x z_L - \kappa^- c_L - k_{cat} c_L \quad (47)$$
$$\dot{w} = \beta - \phi w - \gamma x w \quad (48)$$
$$\dot{y} = k_{cat} c - \phi y \quad (49)$$

with a mass conservation of the total amount of DNA complex constant, $z + c = z^{tot}$ and $z_L + c_L = z_L^{tot}$.

**Steady state analysis.** By making $\dot{c} = \dot{c}_L = 0$, following similar steps as before, we find that $\bar{c} = x z^{tot}/(x + K)$, and $\bar{c}_L = x z_L^{tot}/(x + K)$

$$0 = \theta - \phi \bar{x} - \gamma \bar{x} \bar{w} + (m - 1) k_{cat} z^{tot} \frac{\bar{x}}{\bar{x} + K} + (m_L - 1) k_{cat} z_L^{tot} \frac{\bar{x}}{\bar{x} + K} \quad (50)$$
$$0 = \beta - \phi \bar{w} - \gamma \bar{x} \bar{w} \quad (51)$$
$$0 = k_{cat} z^{tot} \frac{\bar{x}}{\bar{x} + K} - \phi y \quad (52)$$

From equation (50)-(51), we can find $\bar{w}$ as a funtion of $\bar{x}$

$$\bar{w} = \frac{\theta - \phi \bar{x} + (m - 1) k_{cat} z^{tot} \frac{\bar{x}}{\bar{x}+K} + (m_L - 1) k_{cat} z_L^{tot} \frac{\bar{x}}{\bar{x}+K}}{\gamma \bar{x}} = \frac{\beta}{\gamma \bar{x} + \phi}. \quad (53)$$

This leads to a third order polynomial. However, a simple way to find the input-output is to find $\theta$ as a function of $\bar{x}$. This results in

$$\theta(\bar{x}) = \frac{\gamma \bar{x}}{\gamma \bar{x} + \phi} \beta + \phi \bar{x} + \left\{ (1 - m) z^{tot} + (1 - m_L) z_L^{tot} \right\} k_{cat} \frac{\bar{x}}{\bar{x} + K}, \quad (54)$$

and $\bar{y} = k_{cat} z^{tot} \frac{\bar{x}}{\bar{x}+K}$.

**Stability analysis.** We write down the Jacobian matrix for the system (45)-(49), resulting in

$$J = \begin{bmatrix} -\phi - \kappa^+(\bar{z} + \bar{z}_L) - \gamma \bar{w} & \kappa^- + m k_{cat} + k^+ \bar{x} & \kappa^- + m_L k_{cat} + k^+ \bar{x} & -\gamma \bar{x} & 0 \\ \kappa^+ \bar{z} & -\kappa^- - m k_{cat} - k^+ \bar{x} & 0 & 0 & 0 \\ \kappa^+ \bar{z}_L & 0 & -\kappa^- - m_L k_{cat} - k^+ \bar{x} & 0 & 0 \\ -\gamma \bar{w} & 0 & 0 & -\phi - \gamma \bar{x} & \\ 0 & k_{cat} & 0 & 0 & \phi \end{bmatrix}. \quad (55)$$

We inspect the stability of the system numerically by finding the eigenvalues of J. In addition, we can find some conditions for stability when $z_L = 0$, this results in

$$(2\phi + \gamma(\bar{w} + \bar{x}))(\kappa^+\bar{x} + \kappa^- + k_{cat}) + (\phi + \kappa^+\bar{z} + \gamma\bar{w})\phi + (\phi + \kappa^+\bar{z})(\gamma\bar{x}) > (m-1)k_{cat}\kappa^+\bar{z}$$

and

$$\phi(\phi + \gamma(\bar{x} + \bar{w}))(\kappa^+\bar{x} + \kappa^- + k_{cat}) > (m-1)k_{cat}\kappa^+\bar{z}(\phi + \gamma\bar{x}).$$

### 4 Table 1 - General parameters

| Parameter | Value | Annotation | Other studies |
|---|---|---|---|
| $\phi$ (1/s) | $1.2 \times 10^{-3}$ | Degradation RNA | $10^{-4} - 10^{-3}$ [1] |
| $k^+$ (/M/s) | $2.1 \times 10^4$ | STAR and target binding | $10^4 - 10^6$ [2, 3] |
| $k^-$ (1/s) | $6.3 \times 10^3$ | STAR and target unbinding | |
| $k_{cat}$ (1/s) | $0.011$ | STAR-target complex transcription | $0.05 - 0.2$ [4]* |
| $\theta$ (M/s) | $0 - 2.8 \times 10^{-10}$ | STAR-target complex transcription | $2.8 \times 10^{-11} - 2.8 \times 10^{-8}$ [5, 6] |
| $\beta$ (M/s) | $1.4 \times 10^{-10}$ | STAR-target complex transcription | $2.8 \times 10^{-11} - 2.8 \times 10^{-8}$ [5, 6] |
| $\gamma$ (/M/s) | $2.1 * 10^5$ | STAR and sequester binding | $10^4 - 10^6$ [2, 3] |
| $m$ | $5$ | STAR repeats in self-activator | |
| $A$ (M/s) | $2.5 \times 10^{-10}$ | Amplitude of pulse input in self-activator | |
| $T$ (h) | $3$ | Width of pulse input in self-activator | |

* Estimated from the average pause-free velocity of RNAP at saturating concentrations of nucleotide triphosphates (NTP) for an RNA length of $200bp$.

### References

[1] Kim J, Khetarpal I, Sen S, Murray RM. Synthetic circuit for exact adaptation and fold-change detection. Nucleic acids research. 2014;42(9):6078–6089.

[2] Zhang DY, Turberfield AJ, Yurke B, Winfree E. Engineering entropy-driven reactions and networks catalyzed by DNA. Science. 2007;318(5853):1121–1125.

[3] Kim J, White KS, Winfree E. Construction of an in vitro bistable circuit from synthetic transcriptional switches. Molecular systems biology. 2006;2(1):68.

[4] Gabizon R, Lee A, Vahedian-Movahed H, Ebright RH, Bustamante CJ. Pause sequences facilitate entry into long-lived paused states by reducing RNA polymerase transcription rates. Nature communications. 2018;9(1):1–10.

[5] Qian Y, Huang HH, Jiménez JI, Del Vecchio D. Resource competition shapes the response of genetic circuits. ACS synthetic biology. 2017;6(7):1263–1272.

[6] Basu S, Gerchman Y, Collins CH, Arnold FH, Weiss R. A synthetic multicellular system for programmed pattern formation. Nature. 2005;434(7037):1130–1134.
